# Population-scale sequencing resolves correlates and determinants of latent Epstein-Barr Virus infection

**DOI:** 10.1101/2025.07.18.665549

**Authors:** Sherry S. Nyeo, Erin M. Cumming, Oliver S. Burren, Meghana S. Pagadala, Jacob C. Gutierrez, Thahmina A. Ali, Laura C. Kida, Yifan Chen, Fengyuan Hu, Benjamin Hollis, Margarete Fabre, Stewart MacArthur, Quanli Wang, Leif S. Ludwig, Kushal K. Dey, Slavé Petrovski, Ryan S. Dhindsa, Caleb A. Lareau

**Affiliations:** Computational and Systems Biology Program, Memorial Sloan Kettering Cancer Center, New York, NY, USA; Tri-Institutional Program in Computational Biology, Weill Cornell School of Medicine, New York, NY, USA; Centre for Genomics Research, Discovery Sciences, BioPharmaceuticals R&D, AstraZeneca, Cambridge, UK; Department of Radiation Oncology, Memorial Sloan Kettering Cancer Center, New York, NY, USA; Medical Scientist Training Program, Baylor College of Medicine, Houston, TX, USA; Jan and Dan Duncan Neurological Research Institute, Texas Children’s Hospital, Houston, TX, USA; Berlin Institute of Health at Charité – Universitätsmedizin Berlin, Berlin, Germany; Max-Delbrück-Center for Molecular Medicine in the Helmholtz Association (MDC), Berlin Institute for Medical Systems Biology (BIMSB), Berlin, Germany; Department of Pathology and Immunology, Baylor College of Medicine, Houston, TX, USA; Department of Molecular and Human Genetics, Baylor College of Medicine, Houston, TX, USA

## Abstract

Epstein-Barr Virus (EBV) is an endemic herpesvirus implicated in autoimmunity, cancer, and neurological disorders. Though primary infection typically resolves with subclinical symptoms, long-term complications can arise due to immune dysregulation or viral latency, in which EBV DNA is detectable in blood for decades. Despite the ubiquity of this virus, we have an incomplete understanding of the highly variable responses to EBV that range from asymptomatic infection to a trigger for severe disease. Here, we demonstrate that existing whole genome sequencing (WGS) data contains ample non-human DNA sequences to reconstruct a molecular biomarker of latent EBV infection consistent with orthogonal phenotypes, including viral serology. Using the UK Biobank (*n =* 490,560) and All of Us (*n* = 245,394), we uncover reproducible complex trait associations that nominate latent blood-derived EBV DNA as a respiratory, autoimmune, and cardiovascular disease biomarker. Further, we evaluate the genetic determinants of persistent EBV DNA via genome-wide and exome-wide association studies, uncovering protein-altering variants from 147 genes. Single-cell and pathway-scale enrichment analyses implicate variable antigen processing and presentation as a primary genetic determinant of latent EBV persistence, with gene programs expressed in B cells and antigen-presenting cells. Using predicted viral epitope presentation affinities, we implicate genetic variation in MHC class II as a key modulator of EBV DNA persistence. Our analyses demonstrate how existing WGS data can derive novel molecular biomarkers, which may generalize to dozens of viruses comprising the blood virome^1^.

## Main

In 1964, Anthony Epstein, Yvonne Barr, and Burt Achong isolated actively replicating viral particles from Burkitt lymphoma, discovering the virus that would later bear their name – Epstein-Barr Virus (EBV)^2^. EBV has since been identified as the first known human oncogenic virus^3^, the causative agent of infectious mononucleosis^3^, and an agent in developing and exacerbating multiple autoimmune diseases^4^. Despite these wide-ranging pathogenic roles, EBV infection is nearly ubiquitous, infecting >90% of adults worldwide^5^, with most individuals remaining asymptomatic^3^. Further, latent EBV can persist in B cells and even in peripheral blood for the full lifetime of otherwise healthy individuals^1,6^. This clinical heterogeneity – from asymptomatic infection to severe disease – remains incompletely understood. The vast phenotypic spectrum following an endemic infection underscores the individual variability in viral response, which can partially be attributed to genetic variation in the human genome^7–9^. However, genetic association studies of common infections with complex phenotypes such as EBV have been underpowered due to small cohort sizes^10^, motivating new approaches to study infection, viral persistence, and host-phenotype associations.

Beyond its role in human disease, EBV has been instrumental in advancing population genetics research. EBV can transform primary B lymphocytes from healthy individuals into immortalized lymphoblastoid cell lines (LCLs)^11^, enabling long-term storage and large-scale genetic studies, including the HapMap^12^, 1000 Genomes^13^, and Geuvadis^14^ Projects, which applied DNA and RNA sequencing to profile diverse global cohorts. These foundational efforts laid the groundwork for more expansive population-scale cohorts, such as the UK Biobank (UKB^15^) and All of Us (AoU^16^), that include sequencing and phenotypic data from hundreds of thousands of individuals^17^ – a scale that could in theory be used to interrogate the genetic underpinnings of complex, variable phenotypes caused by infection.

As modern biobanks perform whole-genome sequencing (WGS) on peripheral blood rather than LCLs, we hypothesized that latent EBV DNA can be captured and quantified in these existing libraries. Building on recent work that quantifies viral nucleic acids in petabyte-scale datasets to infer host-virus interactions retrospectively^18,19^, we sought to develop a scalable computational heuristic to estimate latent EBV DNA at the level of the individual. By leveraging the existing inclusion of the EBV genome as a contig in the human reference genome, we demonstrate how ordinarily excluded sequencing reads can be reanalyzed to create a novel molecular biomarker for genome-wide and phenome-wide association studies at petabase-scale. Applying this framework to the UKB (*n =* 490,560 individuals; discovery) and AoU (*n* = 245,394; replication) cohorts, we uncover phenotypic and genetic correlates of the variation in latent EBV DNA levels. Our approach reveals the potential for studying genetic determinants of infectious disease across the human virome^20^, which may further serve as a biomarker of other complex traits.

### Biobank WGS data harbors EBV DNA

To address the high levels of EBV DNA present in the LCL-derived libraries that were foundational for efforts such as the 1000 Genomes Project^13^, the EBV genome (chrEBV / NC_007605) was incorporated into the human reference genome assembly (as of hg38)^21^. This alternative contig was designated as a sink for viral nucleic acids to improve variant calling and interpretation in the human genome^21^. We hypothesized that reads mapping to this contig from blood-derived WGS data would reflect prior infection and latent persistence of EBV and serve as a molecular biomarker in these biobank-scale cohorts. We thus extracted all sequencing reads from the aligned.cram files that mapped to chrEBV, enabling a quantification of the per-individual, per-base EBV DNA coverage across 490,560 individuals in UKB (**Fig. 1a,b**). In addition to regions with low coverage corresponding to poor mappability, we identified two distinct loci with disproportionately high read depths that corresponded to repetitive sequences (**Fig. 1b; Methods**). As these regions were detected at levels orders-of-magnitude higher than the median of the viral contig, we reasoned that they would confound latent EBV DNA detection. To assess this, we utilized complementary EBV serostatus as an orthogonal measure of prior infection, evaluated for a subset of 9,964 UKB individuals. Using serostatus, we observed a nominal association between detectable EBV DNA and seropositivity (Fisher’s Exact test odds ratio [OR] = 1.2, *P* = 0.03; **Fig. 1c; Methods**). However, discarding these two repetitive regions revealed that >40% of the UKB cohort only had aligned reads in these regions (**Extended Data Fig. 1b**); masking these regions and re-binarizing the individuals resulted in a markedly stronger association between our bias-corrected DNA detection and serostatus (Fisher’s Exact test OR = 14.6, *P* = 1.7×10^-26^; **Fig. 1c**). Notably, the observed association between EBV DNAemia (here defined as the bias-corrected detection of EBV in blood WGS) and serostatus was specific to EBV (**Fig. 1c,d**). The next strongest association with serostatus was for human immunodeficiency virus 1 (HIV-1; Fisher’s Exact test OR = 4.6, *P* = 0.0023), consistent with previous reports of EBV DNAemia following immunosuppression due to HIV (**Fig. 1d**)^22^. Taken together, these analyses suggest that previously underanalyzed reads mapping to the chrEBV contig represent EBV DNAemia as a novel molecular biomarker. This distinct sequencing-based approach readily scales to hundreds of thousands of individuals, more than a 100-fold increase in sample size compared to serology-based association studies^10^.

**Figure 1.**
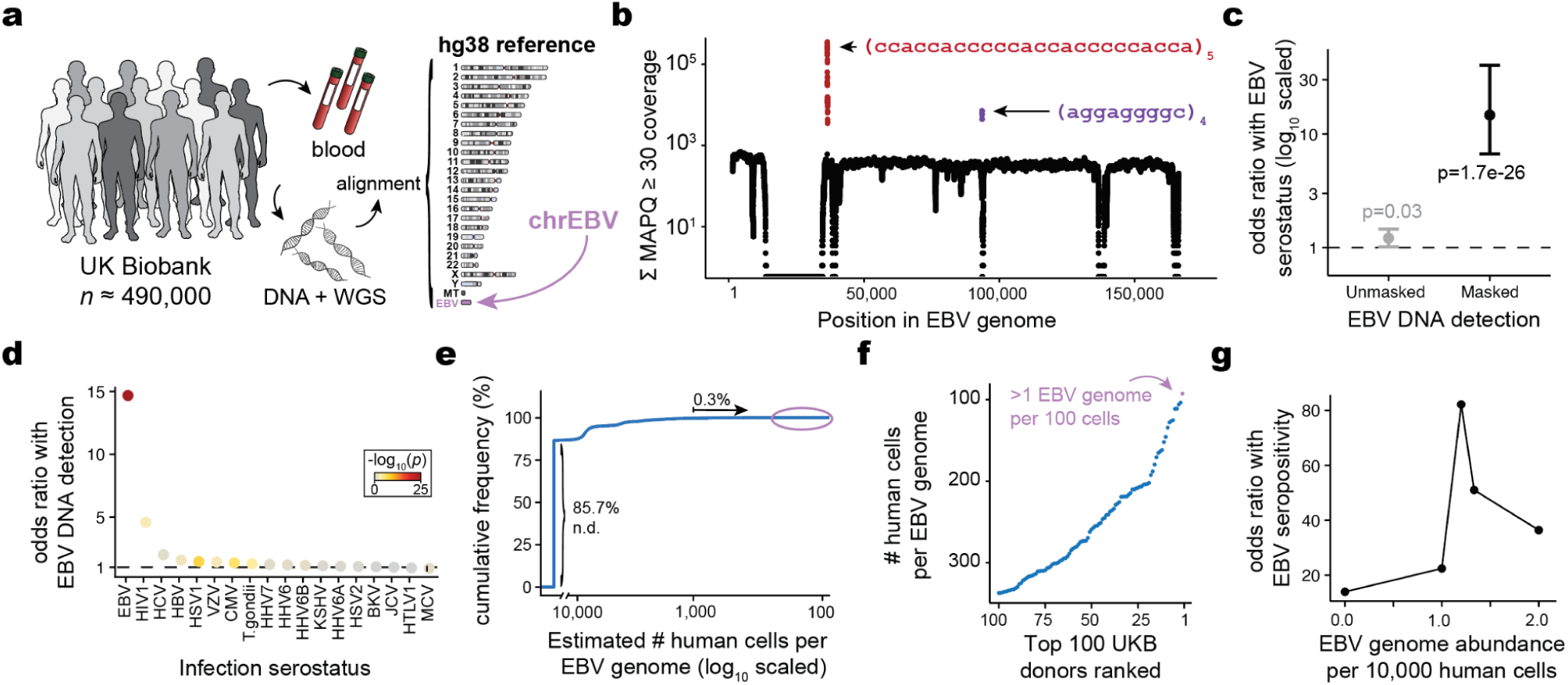
Retrospective quantification of EBV DNA in the UK Biobank. **(a)** Schematic of the approach, in which whole-genome sequencing (WGS) libraries from peripheral blood are aligned to the hg38 reference genome that contains an EBV reference contig (chrEBV). **(b)** Sum of per-base read coverage of high-confidence EBV-mapping reads. Two repetitive regions with inflated coverage are noted in purple and red. **(c)** Association summary of person-level serostatus and EBV DNA quantification with variable region masking. Statistical test: two-sided Fisher’s exact test. Error bars represent 95% confidence intervals for effect estimate. **(d)** Summary of EBV DNA association with serostatus of 18 infectious agents. Statistical test: two-sided Fisher’s exact test. **(e)** Empirical cumulative distribution of detected EBV DNA across the entire cohort. 85.7% of individuals had no detectable (n.d.) EBV DNA. 0.3% had EBV DNA at a copy number of 1+ EBV genome per 1,000 human cells. **(f)** Top 100 individuals based on EBV DNA copy number, from the circled population in (e). **(g)** Association between EBV seropositivity and EBV DNA positivity thresholds at variable levels. Statistical test: two-sided Fisher’s exact test.

To help interpret our metric, we estimated the EBV DNA copy number per 1,000 cells by normalizing read counts between viral and host genome sizes. At the extremes, 85.7% of individuals had no detectable bias-corrected EBV DNA, whereas 0.3% exhibited EBV DNA copy numbers of at least 1 viral genome per 1,000 human cells, including one individual with an EBV genome per 100 human cells (**Fig. 1e,f**). This range, derived from predominantly healthy individuals, is consistent with symptomatic EBV infections being diagnosed at a level of 1 in 200 cells using PCR^23^. Complementary analyses of population-scale scRNA-seq data showed only 1 EBV transcript in >50 billion reads from ∼1,000 blood donors^24^, confirming that the detected viral DNA in UKB is from latent viral infection rather than reactivated or otherwise lytic virus (**Methods**). Given the skewed distribution of EBV DNA load, we assessed varying classification thresholds using serostatus as a ground truth. A cutoff of 1.2 viral genomes per 10^4^ human cells yielded the strongest concordance with seropositivity (OR = 82.2, *P* = 2.2 x 10^-16^; **Fig. 1g**). Hence, using this threshold, we classified 47,452 (9.7%) individuals as EBV DNA+ for subsequent analyses (**Extended Data Fig. 1b**).

Annotating each individual by birth location, we observed an increasing proportion of EBV DNA+ individuals at more northern latitudes in the UK, consistent with prior reports linking EBV infection to vitamin D levels^25^ (**Extended Data Fig. 1c**). Further, we observed a sex-biased (male higher) and age-associated increase in EBV DNA positivity, the latter consistent with EBV serology^10^ (**Extended Data Fig. 1d**). To replicate these inferences, we performed parallel analyses in the US-based All of US (AoU) cohort, spanning 245,394 individuals with available WGS (**Extended Data Fig. 1e; Methods**). Results from the independent analyses of AoU replicated key attributes of the UKB data, including a clear repetitive region that was similarly masked, resulting in 11.9% of individuals annotated as EBV DNA+, as well as consistent age-and sex-associated differences (**Extended Data Fig. 1f–h**). Together, these findings demonstrate that EBV DNA can be retrospectively quantified from existing large-scale WGS datasets with reproducible signal, after mitigating bioinformatic artifacts.

### EBV DNA is a biomarker of complex traits

To determine whether our WGS-enabled measure of EBV DNAemia could serve as a biomarker of complex disease, we performed a phenome-wide association study (PheWAS). Our study used systematic outcomes catalogued via ICD10 codes to identify complex traits highly associated with EBV DNAemia on individuals of predominantly non-Finnish European (NFE) genetic ancestry (**Methods**). Using UKB as a discovery cohort (*n* = 426,563), we tested for the association between EBV DNA+ status and 15,220 binary phenotypes or 1,931 quantitative phenotypes, following our previously described PheWAS workflow^26^ (**Extended Data Table 1; Methods**). Among binary traits, we observed 242 significant (*P* < 3.3 x 10^-6^) associations, including the well-established associations with splenic diseases and Hodgkin lymphoma. We also observed significant associations with rheumatoid arthritis (RA^27^), chronic pulmonary disease (COPD^28^), and systemic lupus erythematosus (SLE^29^), each of which has been previously associated with EBV using orthogonal approaches^6^ (**Fig. 2a**). Significant quantitative associations (*n* = 156) included detection of two EBV antigens, leukocyte count, neutrophil percentage, smoking pack years, and compositions of omega-3 fatty acids, consistent with prior observations of lipogenesis induction following EBV infection^30^ (**Extended Data Fig. 2a**). Interestingly, we also detected a strong association with malaise and fatigue (OR = 1.27; *P* = 2.06 x 10^-10^), noting that EBV has been long hypothesized as a risk factor for myalgic encephalomyelitis/chronic fatigue syndrome (ME/CFS)^31,32^. In addition, we identified significant associations with decreased level phosphatidylcholine (*P* = 2.9 x 10^-9^) and total choline (*P* = 5.9 x 10^-9^), consistent with metabolic studies in ME/CFS patients^33^, reinforcing a potential viral etiology for ME/CFS and suggesting that EBV DNAemia could serve as a facile biomarker.

**Figure 2.**
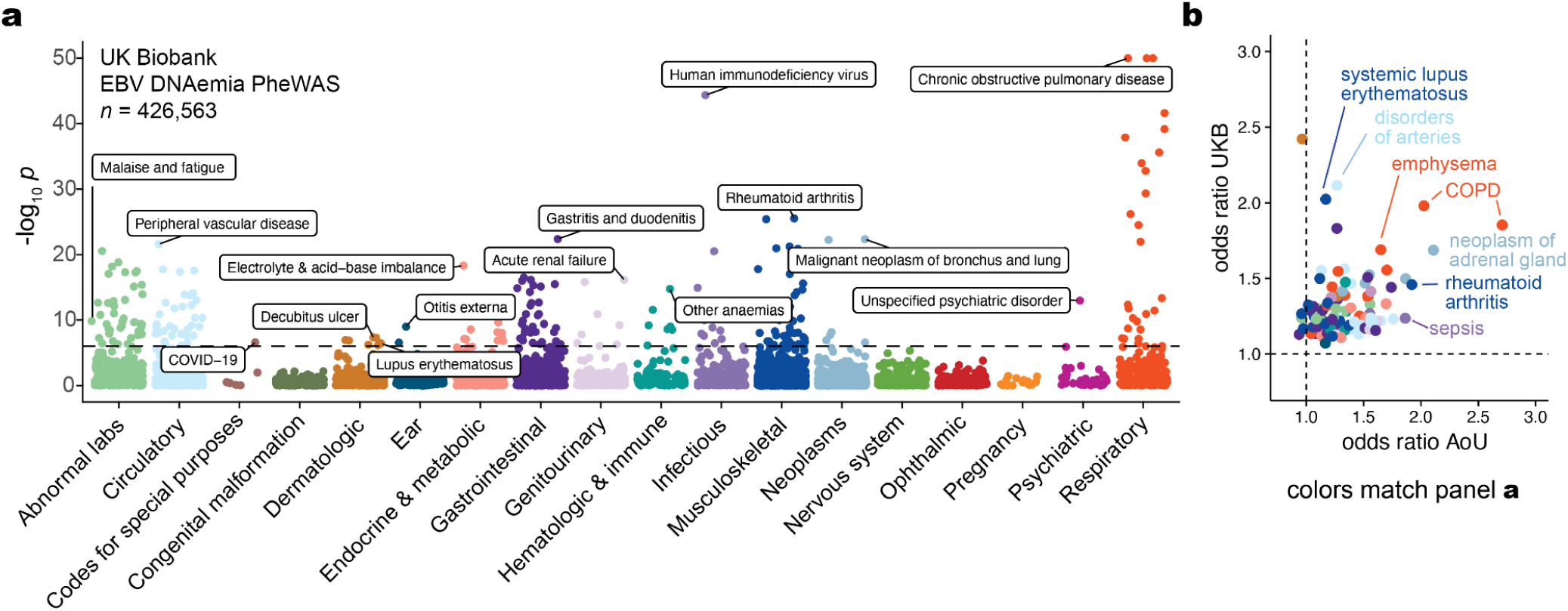
Latent EBV DNA is a biomarker of complex traits. **(a)** Summary of associations between EBV DNAemia and quantitative traits in UKB. The dashed line represents the genome-wide significant *P* value threshold (3.3 x 10^-6^). The y-axis is capped at-log_10_(*P*) = 50; all associations are plotted (*n* = 15,220), and selected traits are highlighted based on biological interest. **(b)** Effect sizes for overlapping ICD10 codes between UKB and AoU. Dotted lines at OR = 1 represent null associations.

We replicated these associations using the AoU cohort (**Fig. 2b**). For 105 of the significantly associated ICD10 codes in UKB, we had sufficient representation in the AoU cohort for replication analyses, 62 of which were also significant in AoU (**Methods; Extended Data Table 2**). These included RA (UKB: OR = 1.46; *P* = 3.9 × 10^-26^; AoU: OR = 1.92; *P* = 6.6 × 10^-20^), COPD (UKB: OR = 1.98; *P* = 1.6 × 10^-33^; AoU: OR = 2.03; *P* = 8.4 × 10^-24^), and lung neoplasms (UKB: OR = 1.50; *P* = 6.7 × 10^-9^; AoU: OR = 1.87; *P* = 0.0062), as well as less-established phenotypes of peripheral vascular disease (UKB: OR = 1.47; *P* = 2.5 × 10^-22^; AoU: OR = 1.39; *P* = 3.1 × 10^-7^), emphysema (UKB: OR = 1.69; *P* = 1.1 × 10^-34^; AoU: OR = 1.65; *P* = 1.1 × 10^-9^), and tachycardia (UKB: OR = 1.30; *P* = 2.2 × 10^-9^; AoU: OR = 1.12; *P* = 0.029), some of which may be attributable to the well-established association between smoking and EBV reactivation^34^. We also considered two additional ICD10 codes for traits previously linked to EBV but were not significant in either cohort (**Methods**; **Extended Data Fig. 2b**). For multiple sclerosis, we observed associations that did not survive multiple testing corrections (UKB: OR = 2.1; *P* = 0.019; AoU: OR = 0.73; *P* = 0.0087), consistent with previous reports that did not observe associations when examining ICD codes of viral exposure on multiple sclerosis outcomes in UKB^35^. For gammaherpesviral mononucleosis, a primary manifestation of EBV infection, the association was consistent in the expected direction (UKB: OR = 2.55; *P* = 0.23; AoU: OR = 5.86; *P* = 1.1 × 10^-6^) but underpowered due to low sample sizes, likely because infectious mononucleosis primarily affects younger individuals (*n* = 11 in UKB; *n* = 42 in AoU). While further work is required to establish a causative role for EBV in previously unreported phenotypes, the concordance of effects across the two biobanks indicates that EBV DNA could serve as a molecular biomarker for these traits, including cardiovascular, respiratory, and autoimmune diseases.

### Polygenic variation underlies EBV DNAemia

Previous studies have established that manifestations of viral infections are a polygenic trait controlled by dozens of loci in the human genome^10,36,37^. Hence, we reasoned that genetic variation would similarly influence the variable detection of EBV DNA. To evaluate this hypothesis, we conducted a genome-wide association study (GWAS) on individuals with primarily NFE ancestry to identify loci associated with EBV DNAemia (**Methods**). Using array-based genotype data followed by imputation from 426,563 NFE individuals in UKB, we identified 21 independent loci reaching genome-wide significance associated with EBV DNA positivity (*P* < 5 × 10⁻⁸; **Fig 3a; Methods**). Conversely, analogous genome-wide analyses of binarized EBV serology (seropositivity) resulted in 0 genome-wide significant hits^38^. We attribute this disparity to both the five-fold increase in sample-size for our NGS-based biomarker, EBV DNAemia, and the underlying molecular biology of the measurements. Namely, seropositivity was measured against four EBV antigens (VCA p18, EBNA-1, ZEBRA, and EA-D) for only ∼2% of NFE participants (n=8,669 out of 426,563 total).

**Figure 3.**
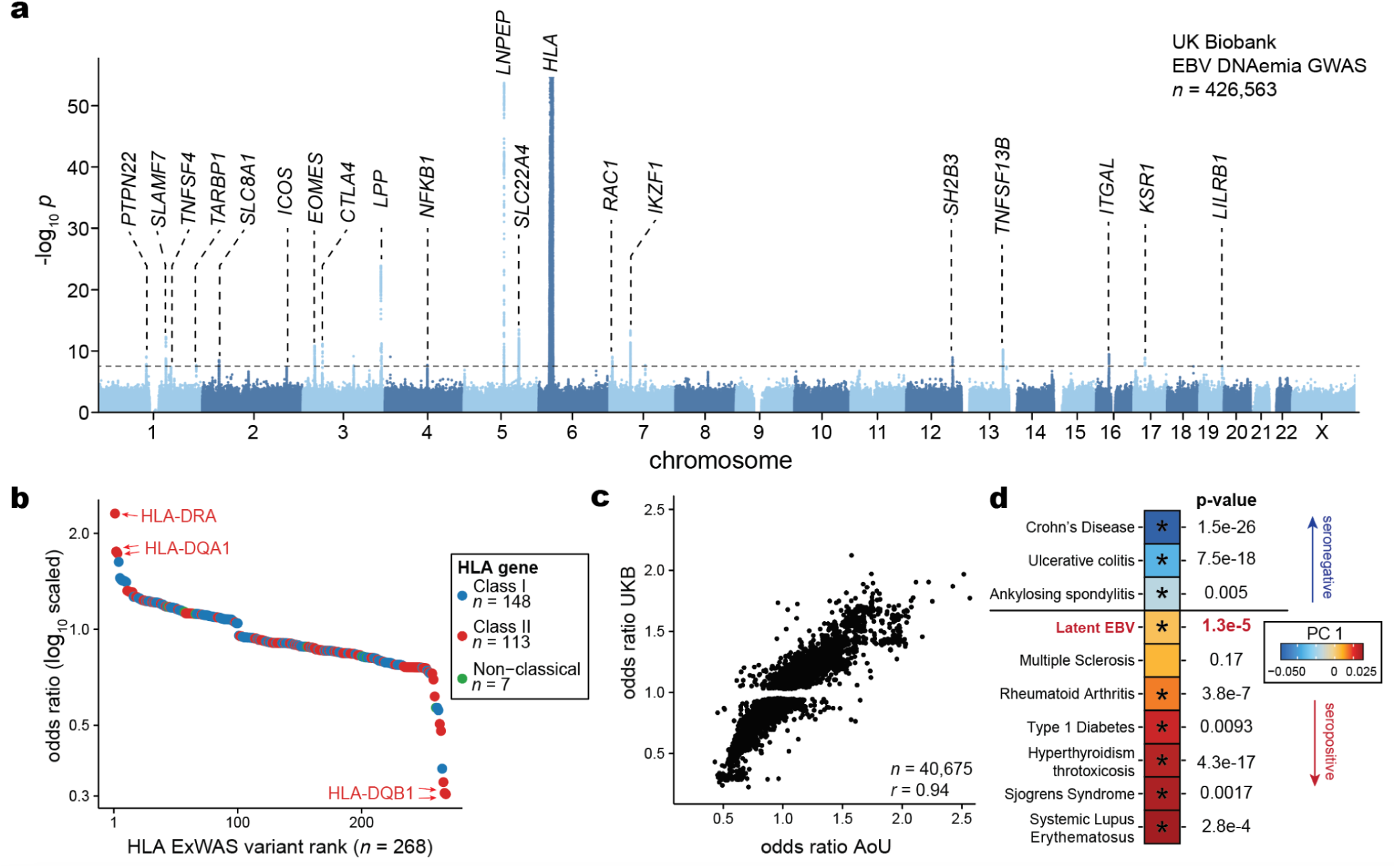
Genetic architecture of EBV DNAemia. **(a)** Manhattan plot summarizing the genome-wide association statistics for EBV DNAemia for 426,563 individuals of predominantly non-Finnish European (NFE) ancestry. Genes proximal to genome-wide significant associations (*P* < 5 × 10⁻⁸) are annotated. **(b)** Summary of protein-altering variants in HLA genes. **(c)** Replication of UK Biobank (UKB)-associated variants in the All of Us (AoU) cohort. The Pearson correlation of variant effect sizes is noted. **(d)** Principal component analysis and projection of EBV summary statistics on complex immune-mediated diseases via cupcake^56^. An asterisk indicates a significant principal component projection score after multiple testing correction.

For the EBV DNAemia GWAS, the strongest association was near human leukocyte antigen (HLA) genes on chromosome 6 that encode the major histocompatibility complex (MHC) class I and II proteins (**Fig. 3a**). MHC molecules are critical in differentiating between self and non-self proteins and have been widely associated with autoimmune traits^10,39,40^. Moreover, associations with viral antibody responses^7–9^ and degree of infection severity^41^ have recurrently implicated this region, corroborating our genetic associations. Overall, the SNP-based heritability (*h^2^*) determined by LDscore Regression (LDSC) was 2.21% (± 0.85%) with limited evidence of genomic inflation (λ_GC_ 1.1; LDSC intercept = 1.03 ± 0.008).

To refine putative functional protein-coding variants at the MHC locus and genome-wide that impact EBV DNAemia, we utilized genome sequencing data from UKB to conduct an exome-wide association study (ExWAS) as we have previously described^26^ (**Methods**). A total of 690 missense variants across 147 genes were significantly associated after Bonferroni corrections (*P* < 5 × 10⁻⁸), which aided in the annotation of putative causal variants at associated loci (**Fig. 3a; Extended Data Table 3; Methods**). Consistent with our GWAS results, the protein-coding variants with the largest effect sizes were in the MHC locus, where 148 MHC class I, 113 MHC class II, and 7 non-classical HLA protein-altering variants were significantly associated with EBV DNAemia (**Fig. 3b**), collectively emphasizing the role of heterogeneous antigen processing and presentation in controlling EBV infection and latent persistence.

Outside the MHC locus, our combined GWAS and ExWAS associations nominated several loci associated with EBV DNAemia. The strongest non-chromosome 6 association was in the chromosome 5 region that encodes the aminopeptidases ERAP1, ERAP2, and LNPEP contiguously^42,43^ (**Extended Data Fig. 3b**). Strongly associated variants included rs2549794 (OR = 0.89, *P* = 3.61 × 10^-51^), an intronic variant in *ERAP2* that modulates gene expression^44^ and manifests with pleiotropic effects on infectious respiratory disease and autoimmunity^44^, consistent with the multi-faceted associations from our phenotypic associations (**Fig. 2**). Additionally, a strong coding variant (rs2476601; R620W) in *PTPN22*, a key immune regulator involved in T cell receptor signaling and type I interferon production^45^, was associated with EBV DNAemia (OR = 1.08, *P* = 1.07 × 10^-9^) (**Extended Data Fig. 3c**). Notably, the *PTPN22* R620W variant has been extensively characterized in autoimmune diseases, including rheumatoid arthritis, systemic lupus erythematosus (SLE), and type 1 diabetes^46^. In addition, this variant has been associated with susceptibility and severity of infectious diseases, including bacterial and viral infections^47^. Moreover, missense variation in *SH2B3* (rs3184504; R262W; OR = 0.96, *P* = 2.43 × 10^-9^; **Extended Data Fig. 3d**) was strongly associated with Celiac disease and linked to type 1 diabetes, peripheral arterial disease, and coronary artery disease, as well as susceptibility to multiple sclerosis^48–52^. Functional studies of this variant (R262W) suggest that *SH2B3* mediates repression of IL12 signaling, promoting enhanced IFN-γ production and hypertension-associated pathology^53^. Other significant non-coding associations with rheumatoid arthritis included rs6679677, located between *RSBN1* and *PHTF1* (OR = 0.93, *P* = 7.41 × 10^-10^), as well as rs3806624 (OR = 0.95, *P* = 6.67 × 10^-11^) and rs9880772 (OR = 0.95, *P* = 1.82 × 10^-11^), positioned in an intergenic region near *LINC01980* and *EOMES*. Notably, both rs3806624 and rs9880772 were also associated with lymphoid malignancies, including Hodgkin’s lymphoma^54^ and chronic lymphocytic leukemia^55^, respectively.

To validate these and other associated loci, we used the AoU cohort as an independent cohort for association analyses to complement the biological plausibility and pleiotropy of genetic associations in UKB. Repeating our GWAS framework on *n =* 131,938 people with EUR ancestry in AoU for 12,099,305 common variants (1% minor allele frequency), we observed largely concordant associations at key loci. Namely, 40,675 variants were genome-wide significant (*P* < 5 × 10⁻⁸) in UKB and passed quality control filters in AoU (**Methods**). Of these, 91.4% were replicated in the AoU GWAS (nominal *P* < 0.05; OR concordant; **Fig. 3c**). These strongly concordant results indicate that persistence of latent EBV DNA is a polygenic trait, and loci underlying EBV DNAemia are reproducible across continents and quantifiable using normally excluded WGS reads. While our discovery and validation methods used the binarized annotation of EBV DNAemia, we note that a linear model considering EBV DNAemia as a quantitative trait yielded very concordant results (**Extended Data Fig. 3e; Methods**).

### Pleiotropy with complex disease

Given the well-described associations between EBV and immune-mediated phenotypes, we sought to systematically evaluate similarities between the genomic architectures of EBV DNAemia and immune-mediated diseases (IMDs). We utilized cupcake^56^, a framework that accounts for the shared components of genetic architecture across 13 IMDs using a shrinkage approach to adjust for LD, sample overlap, allele frequency, and differential sample size via principal component analysis (PCA)^56^ (**Methods**). Using cupcake, we projected the summary statistics from our EBV DNAemia GWAS onto the IMD-derived PCs to assess pleiotropic effects, then tested for significant associations (**Methods**). Notably, the cupcake PC1 segregates an IMD genetic axis characterized by antibody seropositivity^56^. For instance, “seropositive” traits (like RA^27^, SLE^29^, and T1D^57^) have positive PC1, whereas “seronegative” traits (like Crohn’s disease and ulcerative colitis) are negative on this axis (**Fig. 3d**). Consistent with our PheWAS and prior reports of EBV pathogenesis, we observed a positive cupcake PC1 score, reflecting the shared genetic architecture between EBV DNAemia and autoimmune diseases such as RA, SLE, and T1D. Noting that initial EBV infections are most prevalent in adolescence^58^ and generally precede onset of autoimmunity^59^, our data refine a potential model where risk loci to these IMDs may first determine the persistence of latent EBV after primary infection that, in turn, may trigger complications characteristic of disease.

### Cell type-specific and pathway enrichment analyses

To evaluate implicated genes systematically, we examined the expression of the 147 associated ExWAS genes as a signature score in a multi-modal dataset of 211,000 human peripheral blood mononuclear cells (PBMCs; **Fig. 4a**). As expected, the EBV signature score was enriched in B cells, consistent with the known viral tropism of EBV infection and latency (**Fig. 4b,c**). Interestingly, we observed a similar enrichment in subsets of antigen-presenting cells (APCs), particularly conventional dendritic cells (cDCs; **Extended Data Fig. 4a,b**), though DCs are most likely not directly infected by EBV^60^. To resolve the potential biological processes linked to this genetic architecture, we performed gene set analyses (GSA) using the GO Biological Processes (BP) and KEGG Pathway analyses. Among GO BP enriched terms, the top pathways involved antigen processing and presentation, MHC protein complex and assembly, and regulation of T cells (**Fig. 4d; Extended Data Table 4**). From the KEGG enrichments, we observed disease-associated annotations that included viral myocarditis, rheumatoid arthritis (RA), herpes simplex 1 (HSV-1) infection, and, reassuringly, EBV infection (**Extended Data Fig. 4d**). As the strong linkage disequilibrium (LD) on chromosome 6 could drive this association, we refined these enrichments by further removing all HLA-associated genes or all genes on chromosome 6 (**Methods**). Regardless, antigen processing and presentation remained the most enriched term in our GO BP analyses, underscoring the critical role of this pathway in controlling viral infection and clearance (**Fig. 4e,f**). Together, these analyses indicate that B cells and APCs are the primary cell types affected by the genetic architecture of our latent EBV DNAemia biomarker, with viral antigen processing and presentation predominantly influencing the emergence and persistence of latent EBV infection.

**Figure 4.**
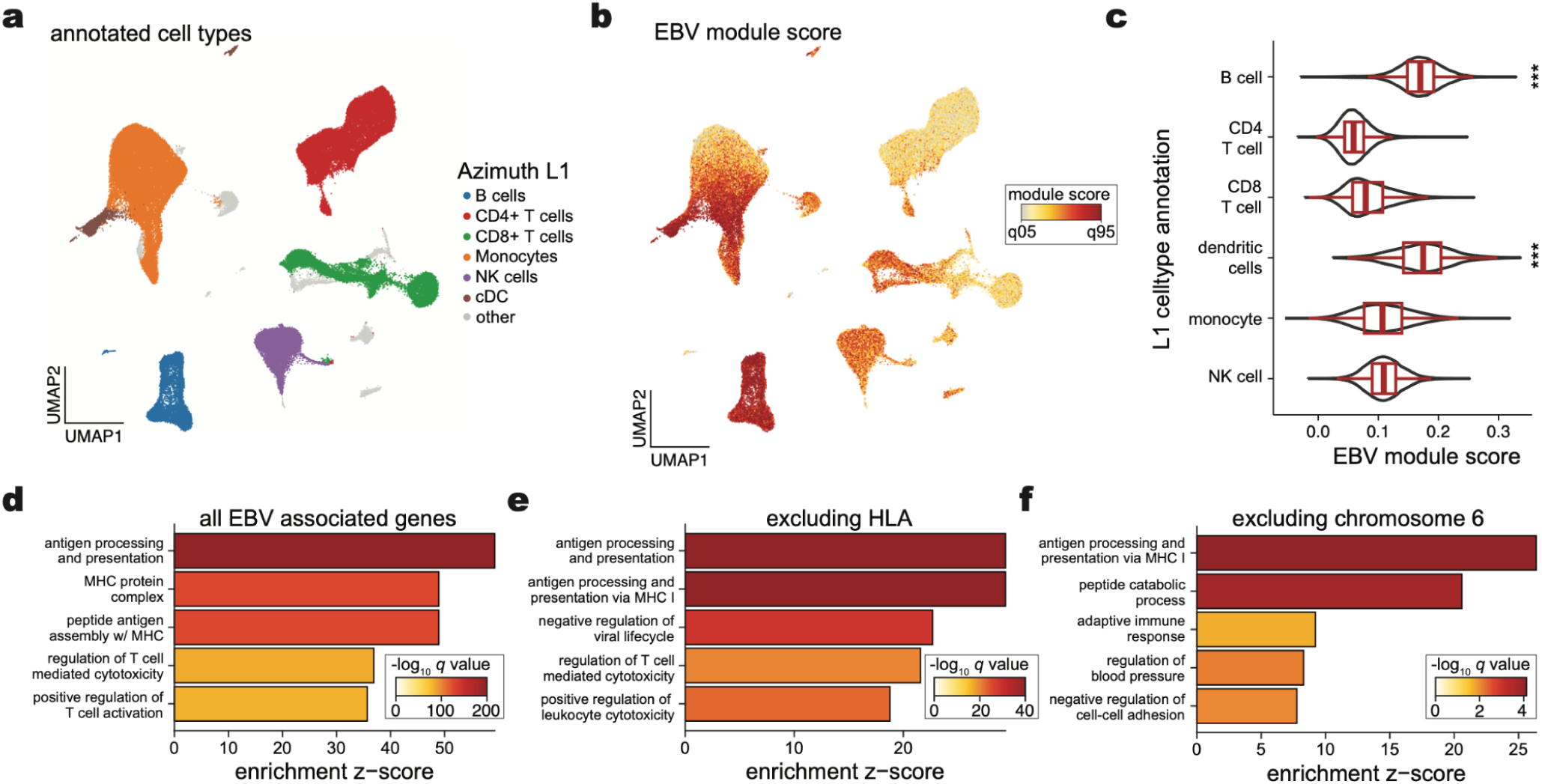
EBV DNAemia gene associations at cell and pathway resolutions. **(a)** UMAP embedding of 211,000 peripheral blood mononuclear cells. The broad cell type annotation (Azimuth L1^73^) annotates major cell types. **(b)** Module score of EBV ExWAS associations, highlighting populations with the highest enrichment. **(c)** Summary of EBV ExWAS scores in major populations. *** indicate statistical significance (*P* < 2.2 x 10^-16^ relative to held-out cell types; one-sided Wilcoxon rank-sum test). Boxplots: center line, median; box limits, first and third quartiles; whiskers, 1.5× interquartile range. **(d)** Summary of top 5 terms from gene set analysis (GSA) of GO Biological Processes Pathways enrichment. **(e)** Same as (d) except excluding annotated HLA genes. **(f)** Same as (d) but excluding all genes mapping to chromosome 6.

### EBV peptide binding strength underlies DNAemia

While the HLA locus is pervasively associated with immune-mediated complex traits, these associations are challenging to resolve due to the allelic diversity of the HLA locus, heterogeneity between human populations, and lack of well-estimated (auto-) antigens that can mediate complex trait manifestation^39^. In our setting, the EBV proteome defines the set of candidate antigens variably presented by these alleles that would, in turn, variably yield EBV DNAemia. Hence, we reasoned that explicit modeling of the HLA variation alongside predictions of EBV peptide display and processing could refine our understanding of genetic variation underlying viral persistence.

We used NetMHC (NetMHCpan and NetMHCIIpan)^61^ to infer the binding affinity of all potential EBV epitopes in the viral proteome with all HLA alleles observed in the UKB NFE cohort (**Fig. 5a; Extended Data Fig. 5a; Methods**). Following prior work that prioritized candidate singular immunodominant epitopes^62,63^, we summarized the per-allele best rank from NetMHC for both class I and class II alleles. The top predicted epitopes prioritized by NetMHC were corroborated by previously identified EBV antigens in the Immune Epitope Database (IEDB)^64^, including 9 of 83 (10.8%) class I peptides and 7 of 106 (6.6%) class II peptides (**Fig. 5b; Extended Data Fig. 5b; Extended Data Table 5**). These overlaps were significantly enriched over a random set of peptides for both class I (*P* = 3.5 × 10^-23^; two-tailed binomial test) and class II (*P* = 0.047; two-tailed binomial test), verifying the capacity for NetMHC to predict viral peptide processing and presentation across HLA alleles in UKB. Further, we observed that predicted immunodominant peptides were depleted in latency-associated EBV genes specifically for MHC class I peptides, reflecting potential viral evolution to evade host immunity during latency (**Fig. 5c**).

**Figure 5.**
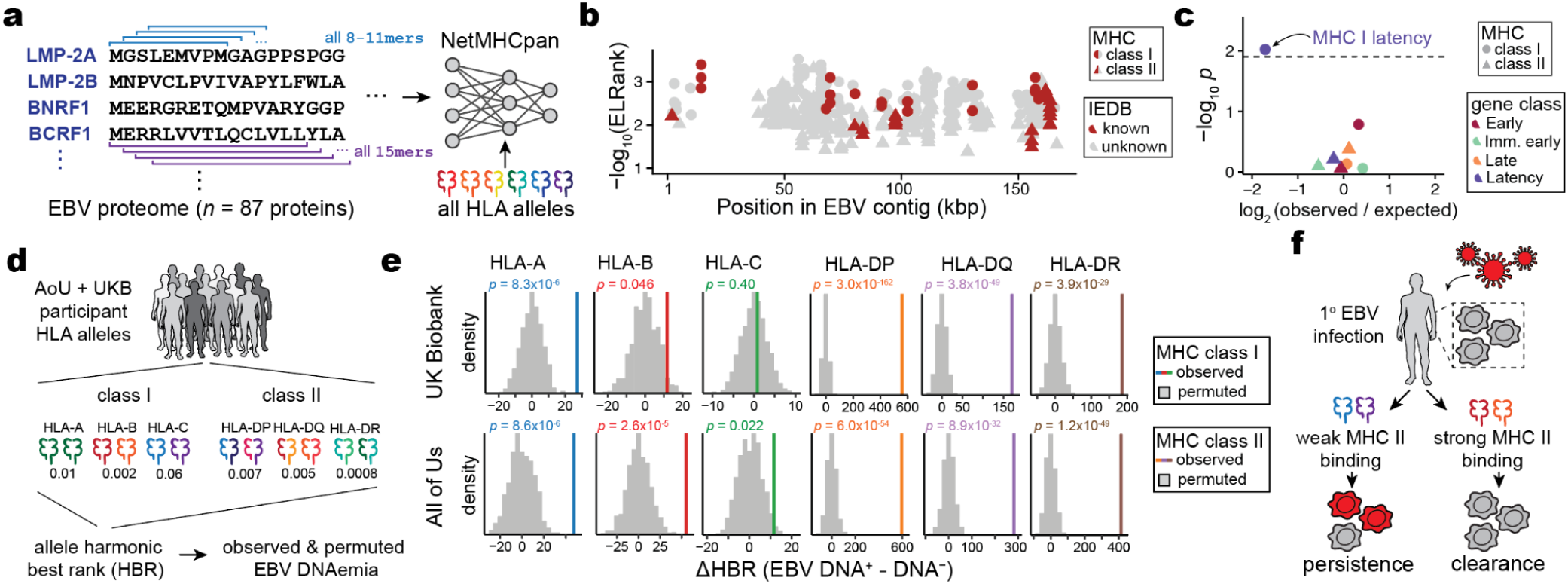
Variable predicted antigen presentation underlies latent EBV DNA persistence. **(a)** Schematic of scoring the EBV proteome for antigen presentation against all HLA alleles observed in the UKB NFE cohort using NetMHCpan and NetMHCIIpan^61^. **(b)** Summary of top antigens bound per HLA class, colored by whether the peptide is annotated in IEDB^64^. **(c)** Enrichment analyses of immunodominant peptides as a function of EBV gene functional class. The highlighted dot shows MHC I peptides are depleted for latency-associated genes. **(d)** Schematic of harmonic best rank (HBR) per HLA allele, which is used as input for downstream analyses. **(e)** Summary of change in comparing individuals with and without detected EBV DNA. *P* values are the result of a permutation test (*n* = 1,000 permutations). **(f)** Overview of an inferred model of antigen processing and presentation via MHC, resulting in persistence or clearance of EBV DNA following infection.

Recent work has shown that aggregation of immunodominant epitopes of the NetMHC scores via a <u>h</u>armonic mean of the <u>b</u>est ranked peptide (HBR) is predictive of immune response, including to neoantigens in tumors^62,63^. We hypothesized that similar measures would be associated with the immune processing and recognition of viral epitopes. Hence, we summarized the per-person, per-allele HBR for class I and class II MHC (**Fig. 5d**). We developed two heuristics to assess the predictive power of these HBR scores in predicting EBV DNAemia, using both a permutation and regression-based framework (**Methods**). We compared the mean difference in HBR for individuals with and without EBV DNAemia and compared against 1,000 permutations of this biomarker. For class I presentation, HLA-A (UKB *P* = 8.3 × 10^-6^) and HLA-B (UKB *P =* 0.046) but not HLA-C (UKB *P* = 0.40) were associated with individual persistence of latent EBV DNA (**Fig. 5e**). Conversely, for class II presentation, each allele was strongly associated (UKB HLA-DP: *P* = 3.0 × 10^-162^; UKB HLA-DQ: *P* = 3.8 × 10^-49^; UKB HLA-DR: *P* = 3.9 × 10^-29^), consistent with the role of CD4-mediated immunity of viral infections via class II antigen presentation by B cells and DCs. These enrichments were concordant with identical analyses in the AoU EUR cohort (**Fig. 5e**). We further verified these results using an additional statistical regression framework that was concordant with our permutation model after accounting for potential confounders, including the full HLA haplotype per individual (**Extended Data Fig. 5c–e; Methods**). Together, these results support a model where individual genetic variation, predominantly in MHC class II, is a key determinant in the latency and persistence of EBV infection (**Fig. 5f**). In particular, our results indicate that the computational modeling between class II host alleles and the viral proteome reflects the development of EBV DNAemia.

### Genetic diversity in EBV sequences

Alongside host genetic variation, our framework enables genome-to-genome analyses^65–67^ whereby variation in the viral genome can be similarly quantified. This setting yields a composition across these biobanks that approximates the circulating genetic variation of EBV in the UK and the US. We developed a heuristic to estimate the ratio of type 1 to type 2 EBV, confirming that type 1 was the predominant strain on both continents^68^ (UKB: 94.8%; AoU: 89.3%; **Extended Data Fig. 6a; Methods**) and indicating the high-quality nature of our data at nucleotide resolution. The overall allele frequencies between the two cohorts were highly correlated (*r* = 0.92), though we observed that UKB detected more EBV variants (*n* = 13,568) than AoU (*n* = 5,471), which could be attributed to the difference in sample sizes or may reflect differences in strain heterogeneity in these two countries (**Extended Data Fig. 6b**).

A longstanding hypothesis is that genetic variation in EBV genomes could explain the diversity in host response ranging from tolerance to pathogenesis^69^, which has been tested by various case-control studies. However, recent reports have also emphasized that variants in EBV previously attributed to oncogenic strains were more closely tied to geographic origin than functional variation^70^. Delineating the potential geographic bias from oncogenic potential is critical, as EBV-driven tumors display stark geographical biases, including nasopharyngeal carcinoma (NPC) that is widespread in southeast China, northern Africa, and other regions in southern Asia^71^. We reasoned that our composite measure of the circulating genetic variation in ostensibly healthy individuals could stratify functional EBV variants of unknown significance (VUS) detected in tumor genomes (**Methods**). Reanalyzing a set of 31 EBV protein-altering mutations from patients with NPC, a tumor driven by EBV infection^70^, we enumerated these VUS based on our observed allele frequencies in UKB and AoU (**Extended Data Fig. 6c**). Notably, all but four variants were detected in one or both cohorts at an allele frequency of 10% (**Extended Data Fig. 6c**). The exceptions were *BALF2* I613V (UKB: 1.7%; AoU: 3.3%), *RPMS* D51N (UKB: 0.19%; AoU: 0.69%), *BNRF1* P694H (UKB: 0.20%; AoU: 0.0%), and *BALF2* V317M (UKB: 0.18%; AoU: 0.0%) (**Extended Data Table 6**). We suggest that the other 27 variants previously detected in NPC genomes are unlikely to be sufficient for pathogenesis, based on their prevalence in healthy individuals in the UK and US. Hence, these VUS likely either reflect geographical drift or require an epistatic effect for driving malignancy^67^. In total, our approach of synthesizing pieces of viral genomes from excluded WGS reads of hundreds of thousands of individuals provides an alternative to low-throughput amplification and sequencing of healthy controls^67,70^ to resolve potential functional variation in the EBV genome.

### Outlook

The exponential rise in population-scale sequencing has transformed our understanding of the genetic determinants of complex phenotypes^17^. While these biobanking efforts were originally genotyped using DNA microarrays, more recent exome and whole-genome sequencing cohorts have discovered a diversity of rare genetic variants underlying complex traits^15,26^. Here, we show that these same large-scale sequencing libraries contain sufficient EBV nucleic acid content to derive novel molecular biomarkers – once corrected for low complexity regions. Our analyses show that host genetic variation underlies the latency and persistence of EBV, with concomitant inferences of the composition of viral heterogeneity across populations. Notably, EBV DNAemia, but not serostatus, is a polygenic biomarker associated with genetic loci impacting antigen processing and presentation machinery and other immune-related signalling. Our framework extends evaluations of endogenous HHV-6 (eHHV-6) that have nominated loci in linkage with germline integration^20,72^, whereas our analyses reveal that latent viral DNA acquired over a lifetime is genetically regulated at a population level. Looking forward, we anticipate that our framework of repurposing existing WGS may elucidate genetic determinants and phenotypic consequences of dozens of viruses and phages that inhabit peripheral blood, which may further resolve the vast phenotypic variation in response to common pathogens.

## Extended Data Tables

**Extended Data Table 1.** Summary of UKB PheWAS.

**Extended Data Table 2.** Reproducible PheWAS associations between UKB and AoU.

**Extended Data Table 3.** Summary of significant genes and variant associations from ExWAS.

**Extended Data Table 4.** Full results of pathway enrichment analyses.

**Extended Data Table 5.** Annotation of immunodominant epitopes per HLA alleles with IEDB annotation.

**Extended Data Table 6.** Full annotation of population allele frequencies for EBV variants of unknown significance.

## Methods

### Rationale of EBV detection

The 171,823 nucleotide EBV genome (NC_007605.1) was first included in December 2013 (hg38 version GCA_000001405.15) as a sink for off-target reads that are often present in sequencing libraries, to account for pervasive EBV reads present from the immortalization of LCLs (as with the 1000 Genomes Project and related consortia). Importantly, whole-genome sequencing (WGS) in the UK Biobank (UKB) and All of Us (AoU) consortia was performed on whole blood or whole blood subfractions^16,74^, reflecting that EBV reads detected would derive from latent viral DNA from prior infections.

As an independent measure of validating that EBV is latent (rather than an active infection), we quantified scRNA-seq data from peripheral blood mononuclear cells from ∼1,000 individuals profiled via single-cell sequencing^24^ using kallisto^75^ for mRNA quantification (analogous to our previous execution with HHV-6^18^). We reanalyzed a total of 53,872,337,003 paired-end sequencing reads from this consortium, identifying only 1 UMI that was classified as uniquely mapping to the EBV transcriptome (the BARF0 EBV gene). Hence, our interpretation of the detection of EBV DNA from WGS libraries is a measure of residual latent virus that is not active for nearly all individuals. As ∼90% of individuals (in UKB and in these populations) are seropositive, yet we detect high-confidence EBV DNAemia in ∼10% of individuals, we interpret our measure to reflect the tail of latent viral retention among individuals for whom this is highest.

### WGS data and cohort analyses in UKB

For UKB, we obtained per-base abundance of EBV DNA of the 490,560 WGS libraries by extracting reads aligning to chrEBV in the hg38 human genome reference with a read mapping quality (MAPQ) ≥30 via the samtools view command^76^. To quantify EBV DNA abundance for each position, we summed the coverage of each base in the EBV genome across all libraries (per-base abundance). The resulting coverage across the viral contig was approximately flat, supporting that EBV DNA detection from WGS reads was real viral DNA, with two key exceptions (**Fig. 1b**). First, a total of 27,692 positions had low-to no coverage (per-base abundance ≤ 10) due to low mappability of the EBV contig. Second, two regions (positions 36,390-36,514 and 95,997-96,037) had orders-of-magnitude higher coverage (per-base abundance ≥ 10^3^ at these 166 positions). Upon further examination, the sequences were highly repetitive (**Fig. 1b**). Hence, we reasoned that these two regions may confound EBV DNA detection. To assess this, we calculated EBV DNA abundance per person before and after masking, by summing MAPQ ≥ 30 coverage either across all *J* = 171,823 bases, or only across the remaining *J’* = 143,965 well-covered bases (10 < per-base abundance < 10^3^ for each base). The per-individual EBV sum unmasked was computed over all *J* bases, whereas the masking was performed over *J’* bases.

We then used a two-sided Fisher’s exact test to test for association between EBV DNA presence (EBV DNA coverage > 0) and EBV serostatus (“EBV seropositivity with Epstein-Barr virus,” a measurement in UKB with ID 35811742). Before masking, EBV DNA presence had a weak but insignificant positive association with EBV seropositivity (OR = 1.2, *P* = 0.03). Conversely, after masking these repetitive regions and recomputing donor positivity, the association between DNAemia and seropositivity was much stronger (OR = 14.6, *P* = 1.7 x 10^-26^) (**Fig. 1c**). These analyses demonstrate that masking highly-repetitive regions in the viral contig are required to perform valid inferences of latent viral activity retrospectively. In other words, we highlight that a simple enumeration of aligned reads is insufficient to uncover high-quality associations, and further processing is required for low-complexity regions, as evidenced by statistical overlap with EBV serostatus.

### Contig mappability analyses

To confirm that regions of the EBV contig that were not detected were attributable to poor mapping quality of those regions, we generated synthetic reads of length 101 bases by tiling the reference EBV contig. Next, each synthetic read was aligned using bowtie2^77^. We define mappability as the percentage of reads overlapping a position with a map quality score exceeding 10. This analysis reproduced regions depleted from the pseudobulk abundance (**Extended Data Fig. 1a**), indicating that lowly detected regions were due to homology in the hg38 reference rather than variable DNA presence in the latent EBV genome.

### EBV DNA copy number estimation and thresholding

To calculate EBV DNA abundance per person, we summed the coverage over the well-covered, non-biased bases (*J’*). We normalized this value against the effective EBV genome size (143,965 bases) to get an estimate of the coverage per EBV genome. Next, we used the human WGS coverage and accounted for the diploid human genome to compute an estimate of EBV DNA copy number per human cell. The resulting value was an estimate of the EBV DNA copy number per cell, which was predominantly 1 in 1,000-10,000 cells for individuals whom this was detected (i.e, our limit of detection was approximately 1 EBV genome per 10,000 cells). To the best of our knowledge, analogous values have not been comprehensively computed in healthy populations, but clinical diagnostics of EBV infection estimate loads of 1 in 200 cells using PCR for positive diagnoses^23^. While a previous study similarly used EBV reads in a cohort of ∼8,000 donors, this analysis used reads that did not map to the human reference genome and did not correct for the repetitive, biased DNA abundances that significantly skewed the resulting quantification^1^. After quantifying per-person EBV DNA abundance, 85.7% of individuals in UKB had no detectable EBV DNA. Further, we noted a small fraction of individuals (*n =* 365) with minimal EBV DNA (i.e., only 1 high-confidence base after excluding repetitive regions). The extreme left skew of this distribution suggests that converting EBV DNA copy number to a binary trait, EBV DNA detection, would be more suitable, since a quantitative trait otherwise assumes a dose-dependent relationship when testing for associations.

Using a two-sided Fisher’s exact test, we surveyed different cutoffs against association with EBV serostatus to determine an optimal EBV copy number threshold. We observed the most significant positive association with a threshold of 1.2 EBV copies / 10^4^ human cells (OR = 82.17, *P* ≈ 0). This corresponded to having a per-person abundance of at least 302 bases covered, which corresponds to a full paired-end sequencing read (2 x 151bp) with no soft-clipping. 47,452 people (9.67%) had EBV copy numbers greater than this threshold, which was used for all downstream analyses.

### EBV DNA detection in AoU

We obtained per-base abundance of EBV DNA for 245,394 people in AoU with WGS data similarly by extracting reads that mapped to chrEBV in the hg38 human genome reference with an alignment quality (MAPQ score) of ≥ 30. To quantify EBV DNA abundance per base, we summed the q30 coverage of each base in the 171,823 bp EBV genome across all people. We again observed an overall uniform coverage; 23,513 positions had no coverage (per-base abundance = 0), and 4 regions (positions 36,389-36,516; 52,012-52,034; 95,997-96,037; and 163,596-163,617) had abnormally high coverage (per-base abundance >1,000 at 214 positions; **Extended Data Fig. 1g**). The effective EBV genome size was the remaining 148,096 bases (> 0 but < 10^3^ for each base). While the largest repetitive region was the same in both UKB and AoU, differences in the other regions with variable bias could be attributed to differences in the alignment software for either cohort, noting that all analyses used the existing mappings from either cohort.

We quantified the EBV copy number per person in AoU with a similar approach as used in UKB. Briefly, we normalized by the effective EBV genome size, then by the average genome coverage (30x human WGS) provided by AoU metadata. A total of 51,459 people (21%) had detectable EBV DNA. The top EBV DNA load harbored was ∼1 EBV copy per 1.4 cells (or 7,046 EBV copies per 10^4^ cells) (**Fig. 1c,d**). Using the same EBV DNA copy number thresholds as in UKB, a total of 29,249 people (11.9%) had EBV copy numbers greater than the threshold of 1.2 EBV copies per 10^4^ human cells. The overall higher EBV loads in AoU compared to UKB may be due to a difference in the recruitment criteria and demographics of the two cohorts: relative to the general population (as in AoU), UKB shows a “healthy volunteer bias” where participants were less likely to have self-reported health conditions^17^. In comparison, the maximum described in a previous paper was a few orders of magnitude higher (2,404,531 EBV copies per 10^5^ human cells), potentially due to our exclusion of abnormally high coverage regions^1^.

### Phenome-wide association studies

We conducted PheWAS in UKB as a discovery cohort to test for the association between EBV DNAemia and 13,289 binary phenotypes and 1,931 quantitative phenotypes amongst participants with broadly non-Finnish European ancestry (NFE) as in the GWAS (refer to the following section). We employed logistic regression with Firth correction and included the same covariates as in the GWAS: age + sex + age * sex + age^2^ + age^2^ * sex + batch + ancestry PCs 1-20. Using a Bonferroni correction, we defined 0.05/15,220 = 3.3×10^-6^ as our significance threshold.

We replicated PheWAS associations using the AoU cohort via Fisher’s exact tests for association between EBV DNAemia and each representative ICD9/10CM code in AoU. As recommended in the AoU workbench, we defined a representative ICD code as a code appearing at least twice in a person and 20 instances across all participants. The top results were predominantly being HIV positive, having immunodeficiencies, or receiving organ transplants, which we also observed in UKB. To compare effect sizes between hits in UKB and AoU, we matched AoU ICD10CM codes to a corresponding ICD10 code by taking the first four characters of the ICD10CM code, as codes > 4 characters do not exist in the ICD10 ontology used in UKB.

### Genetic associations with EBV DNAemia in UKB

For individuals with broadly non-Finnish European (NFE) ancestry in UKB, array-based imputed genotypes with good genome-wide coverage in the common (>5%) and low frequency (1–5%) MAF ranges were available^15^. Genotyping arrays capture genome-wide genetic variations (SNPs and indels) within both coding and noncoding regions, allowing imputation of genotypes and tests for association between genotypes and a specified trait. To avoid confounding results due to differences in ancestral background, we stratified the cohort across six broad genetic ancestries (AFR, AMR, ASJ, EAS, NFE, and SAS) before testing for associations between EBV DNAemia and UKB-imputed genotypes, which resulted in a total of 450,032 individuals with array imputed genotype data available, including 426,563 individuals of non-Finnish European (NFE) ancestry. We then used REGENIE v3.5^78^ to examine associations between EBV DNAemia (EBV DNA+ status) and imputed genotypes, using a logistic model with covariates and applying Firth correction: EBV DNAemia ∼ age + sex + age * sex + age^2^ + age^2^ * sex + batch + ancestry PCs 1-20, as previously described. The input to REGENIE includes directly genotyped variants (MAF>1%, MAC>100, genotyping rate per variant >99%, and genotyping rate per individual >80%). We pruned these variant sets using PLINK2 (--indep-pairwise 1000 100 0.8) as input to REGENIE’s step1 analyses. This step produces a whole genome regression model to fit to the binary trait of EBV DNAemia and outputs a set of genomic predictions.

For REGENIE step2, we further filtered out SNPs that had 0.99 “missingness,” imputation INFO < 0.7, and p.HWE > 1 x 10^-5^. This step fits a logistic model to imputed data, using the genomic predictions from step1. To estimate heritability of SNPs and genomic inflation, we performed LD score regression (LDSC) by applying the ldsc package (v1.0.1). Briefly, we used munge_stats.py on the cleaned summary stats, then used ldsc.py to estimate h^2^ using the supplied 1KG Genomes LD score matrices. Identical steps were applied to execute the EBV serology on the subset of patients where multiplexed serostatus was measured^38^ (**Extended Data Fig. 3a**).

To annotate variant loci, we focused on significant variants (*P* < 5 x 10^-8^) and created genomic intervals of ± 1 Mb around each variant. As variants on chromosome 6 often exhibit linkage disequilibrium with MHC, we created a custom interval (chr6: 25,500,000 to 34,000,000) for the HLA region. We then combined overlapping intervals using the GenomicRanges *reduce* function and selected the most significant variant per interval as the index variant. In the case of ties, we selected the variant closest to the midpoint of the region. We applied the *reduce* function again to ensure we had a set of non-redundant index variants. Finally, we annotated each variant by the closest gene, using the Bioconductor library biomaRt to access Ensembl v111 (Jan 2024) gene annotations and selecting the gene whose midpoint was closest to the index variant. For visualization of specific loci, we used the canonical hg38 reference genome isoforms. Linkage disequilibrium was determined via LDlink^79^ for the regions noted (**Extended Data Fig. 3b-d**). Zoom plots were from the array-based GWAS associations in UKB, and the LD reference panel in LDLink^79^ used all EUR populations.

We complemented our GWAS with an exome-wide association analysis (ExWAS), leveraging the whole-genome sequencing data available in UKB. Specifically, we tested for associations between EBV DNAemia and protein-coding variants and observed at least six participants of European ancestry in UKB. We applied our previously described protocol to generate variant-level statistics^26,80^. Variants were required to pass the following quality control (QC) criteria: coverage ≥10x; ≥0.20 of reads with the alternate allele for heterozygous genotype calls; binomial test of alternate allele proportion departure from 50% in heterozygous state *P* ≥ 1 × 10−6; GQ ≥ 20; Fisher Strand Bias (FS) ≤200 for indels and ≤60 for SNVs; root-mean-square mapping quality (MQ) ≥40; QUAL ≥30; read position rank sum score (RPRS) ≥−2; mapping quality rank score (MQRS) ≥−8; DRAGEN variant status = PASS; and ≤10% of the cohort with missing genotypes. Additional out-of-sample QC filters were also imposed based on the gnomAD v2.1.1 exomes (GRCh38 liftover) dataset^81^. The sites of all variants were required to have ≥10x coverage in ≥30% of gnomAD exomes and, if present, each variant was required to have an allele count ≥50% of the raw allele count. Variants with missing values for any filter were retained unless they failed another metric. Variants failing QC in >20,000 people were also removed. *P* values were generated via Fisher’s exact two-sided test. Three distinct genetic models were studied for binary traits: allelic (A versus B allele), dominant (AA + AB versus BB), and recessive (AA versus AB + BB), where A denotes the alternative allele and B denotes the reference allele.

### Replication of UKB EBV DNAemia-associated genotypes in AoU

To broadly capture variants in individuals with imputed European ancestry within AoU, we utilized the variant-level metadata for the SNP and indel variants contained in the short read WGS (srWGS) data dictionary. We filtered for variants with an alternative allele frequency (AF) of 0.01 < AF < 0.49 or 0.51 < AF < 0.99 (gvs_eur_af) and at least 100 individuals containing this variant (gvs_eur_sc ≥ 100) in the European subpopulation as the input variants lists to step1 and 2 of the REGENIE pipeline. This resulted in 16,566,413 variants across chromosomes 1-22. EBV DNAemia was supplied as a binary trait, along with the covariates age, sex, age*sex, and ancestry PCs 1-15. There were 133,578 such individuals that had EBV DNAemia levels determined, of which 131,938 had complete covariate data and were used in the analysis, along with 12,099,305 total variants.

### Genomic architecture associations

To holistically evaluate genetic architecture similarities between EBV DNAemia and IMDs, we used the R package cupcake. The package was used to define shared components of genetic architecture across 13 IMDs, applying shrinkage to adjust for LD, individual overlap, allele frequency, and differential sample size. Summary statistics of 13 large IMD GWAS were projected into a reduced dimension space, which served as a common genetic basis that enabled simultaneous comparisons between multiple diseases. A set of 566 variants was extracted, and PCA was applied to distill the variants down into 13 PCs that were defined as genetic risk components^56^. Applying this approach, we first used tabix to extract summary association statistics for these 566 variants from our NFE EBV status GWAS. After checking and adjusting the effect allele alignment, we used cupcake^56^ to project these variants onto the 13 IMD genetic risk components and assess the significance of association with each component.

### Pathway and single-cell analyses

To evaluate the gene expression program uncovered by our ExWAS association, we utilized a high-resolution single-cell CITE-seq dataset of peripheral blood mononuclear cells from eight distinct donors with 210,911 quality-controlled cells^73^. The 147 ExWAS-associated genes were input alongside the preprocessed Seurat object into the AddModuleScore function with default hyperparameters. We removed genes mapping to the HLA region as well as ribosome-associated genes from the input gene list due to technical variation (HLA: genetic polymorphisms; ribosome: cell quality) from the module score foreground and background. Downstream association analyses of cell type enrichment were performed using the pre-supplied labels.

Pathway enrichment analyses were performed using the same ExWAS gene set via the clusterProfiler R package^82^. Gene set analyses were performed using the enrichGO (for biological processes) and enrichKEGG functions using the set of 147 genes and all ENSEMBL human genes as a background set. For analyses with HLA (**Fig. 4e**) and chromosome 6 excluded (**Fig. 4f**), we removed appropriate genes both from the foreground (i.e., test set) and background set for statistical analyses. We used the simplify() function in clusterProfiler with a similarity cutoff of 0.7 (the default value) to reduce the number of redundant association terms. Hence, we note that the labels in panels **Fig. 4d-f** are not identical in name; this result is due to the simplify() function’s selection of a single term that is nearly identical to other related terms.

### HLA haplotype and EBV peptide presentation

We used the four-digit HLA imputation calls processed in the UK BiobankRAP that were called via HLA*IMP:02^83^. In brief, allele dosage values >0.7 were used for defining donor haplotypes for a specific four-digit HLA allele. Homozygotes were determined by alleles with values >1.3. For All of Us, predetermined HLA genotypes were not available in the workbench. Hence, we reconstructed the HLA calls for individuals of EUR ancestry, using the T1K toolkit using a synthesis of chromosome 6 mapping reads, reads mapping to HLA decoy contigs, and unmapped reads^84^. Following T1K toolkit recommendations, the donor haplotypes were determined by alleles with a Quality score >0. In addition, homozygotes were determined by donors with only a single allele called and with a Quality score >30.

The amino acid sequences of all 87 unique EBV protein sequences were obtained from the peptide sequence of the nuccore NC_007605. The protein fasta file was input to NetMHCpan, along with all observed MHC class I alleles (HLA-A, HLA-B, or HLA-C) and class II alleles (HLA-DR, HLA-DP, or HLA-DQ) in the UKB NFE cohort. Sliding windows of all 8-, 9-, 10-, or 11-mers of the provided protein sequences were generated for prediction of class I allele peptide presentation, and sliding windows of size 15-mers for class II. These peptides were scored for available alleles that could be quantified via NetMHCpan4.1 and NetMHCIIpan4.3^85^.

The NetMHC output reflects the predicted %rank score for each peptide and a given allele, which is a measure of the rank of the predicted affinity of the allele for the peptide compared to a set of 400K random natural peptides. For MHC class I, we computed the harmonic best rank (HBR) score per allele by taking the harmonic mean over the two genotyped alleles for each of HLA-A, B, and C. For homozygotes, the harmonic mean is equivalent to any individual observation. For individuals missing a single allele, we considered only the genotyped call, and for two missing alleles, the individual was excluded from the per-allele analysis.

For MHC class II analyses, all HLA-DRB alleles were directly applied as input, along with the EBV proteome fasta file, to generate all possible 15-mer sliding windows. As HLA-DQ and HLA-DR alleles exist in pairs of alpha and beta alleles within the predictions, we took all HLA-DQ and HLA-DP alleles imputed in the UKB NFE cohort and generated all possible combinations of HLA-DQA/HLA-DQB alleles and all possible combinations of HLA-DPA/HLA-DPB allele pairs. These alpha-beta allele combinations were then used as inputs to NetMHCIIpan, along with the EBV proteome fasta file. Again, the output file lists each peptide, the protein from which the peptide is derived, a given class II allele (pair), and the predicted %rank_EL score, which is the percentile rank of the eluted ligand prediction score. As HLA-DRA is the only non-variable gene in the population, each individual has only two possible HLA-DR heterodimers. Each individual can form four possible alpha-beta heterodimers from HLA-DP and HLA-DQ (between alpha and beta molecules). Hence, each individual may assemble 10 possible unique heterodimeric MHC-II molecules^62^.

The per-allele HBR was computed using the harmonic rank of the heterodimers for each allele class and rescaled by a factor of 10^6^ when computing the final ΔHBR score (shown in **Fig. 5**). The comparisons were only between the NFE (EUR) populations in either cohort identified through genetic analyses. To further verify our effect was linked to class II presentation strength, we completed regression analyses using the same set of covariates for our genetic association analyses, which verified that other forms of confounding (e.g., population stratification; sex) did not explain the associations between the class II predicted presentation strength and EBV DNAemia.

### EBV viral sequence analysis

Raw sequencing reads from chrEBV were merged from all participants from both cohorts. The aggregated.bam file was transformed into a per-base, per-nucleotide count using bam-readcount^86^. For the type 1 and type 2 strain analyses, we sought to quantify the abundance directly from the aligned reads to the chrEBV reference (a type 1 EBV strain). Here, we performed a multiple-sequence alignment of the EBNA-2 gene (the major difference between strains) for nuccore IDs K03333 (type1) and K03332 (type 2) and mapped the MSA coordinates back to the chrEBV reference to identify putative regions that would reflect single nucleotide variation that would, in turn, reflect strain-level differences. We identified nine variants (all on chrEBV): 36209C>T, 36226T>A, 36251A>G, 36252A>T, 36258C>A, 36275G>T, 36302A>C, 36312T>A, and 36320C>T, where the reference was type 1-derived and the alternate was type 2-derived. These were selected based on: (a) no more than 1% allele frequency aside from the ref and most abundant alt; (b) no overlap with the repetitive regions (**Fig. 1b**); and (c) filtered for overall consistency. A set of mutations of 31 protein-altering mutations in EBV (**Extended Data Fig. 6c**) was curated from a recent resource and global-scale analyses of EBV genomes^70^ derived from individuals with EBV+ nasopharyngeal carcinomas.

## Acknowledgements

We are grateful to the participants and staff of the UK Biobank and US-based All of Us for enabling the research. We thank T. Rückert, C. Honeycutt, T. Nawy, and members of the Lareau and Dhindsa labs for their helpful feedback. This work was directly supported by the Human Virome Program (U01AT012984 to C.A.L. and R.S.D.), P30CA008748 (to C.A.L. and K.K.D), and a Michelson Prize Next-Generation Grant (C.A.L.). C.A.L is supported by R00HG012579 and a Frank Howard Scholar Award. K.K.D. is supported by R00HG012203 and R01HG014008.

## Author contributions

S.S.N., S.P., R.D., and C.A.L. conceived and designed the study. S.S.N., E.M.C., O.S.B., R.D., C.A.L. led bioinformatics analyses. M.S.P., J.C.G., T.A., L.C.K., F.H., B.H., M.F., S.M., and Q.W. supported the bioinformatics analyses. K.K.D. and L.S.L. aided in data interpretation. S.S.N., E.M.C., O.S.B., S.P., R.D., and C.A.L. wrote the manuscript with input from all authors.

## Code Availability

Code to reproduce custom analyses in this manuscript is available online at https://github.com/clareaulab/ebv_biobank_gwas.

## Data Availability

The UK Biobank data are available to qualified researchers (please refer to the details at http://www.ukbiobank.ac.uk/register-apply/). The All of Us data are available as a featured workspace to registered researchers of the All of Us Researcher Workbench (https://www.researchallofus.org/). Summary statistics from the EBV DNAemia discovery GWAS in UKB is available at https://my.locuszoom.org/gwas/409414/?token=6385c90400414f34b8ed17679bf1495b and have been uploaded to the GWAS catalogue (GCST90572743).

---

**Extended Data Figure 1.**
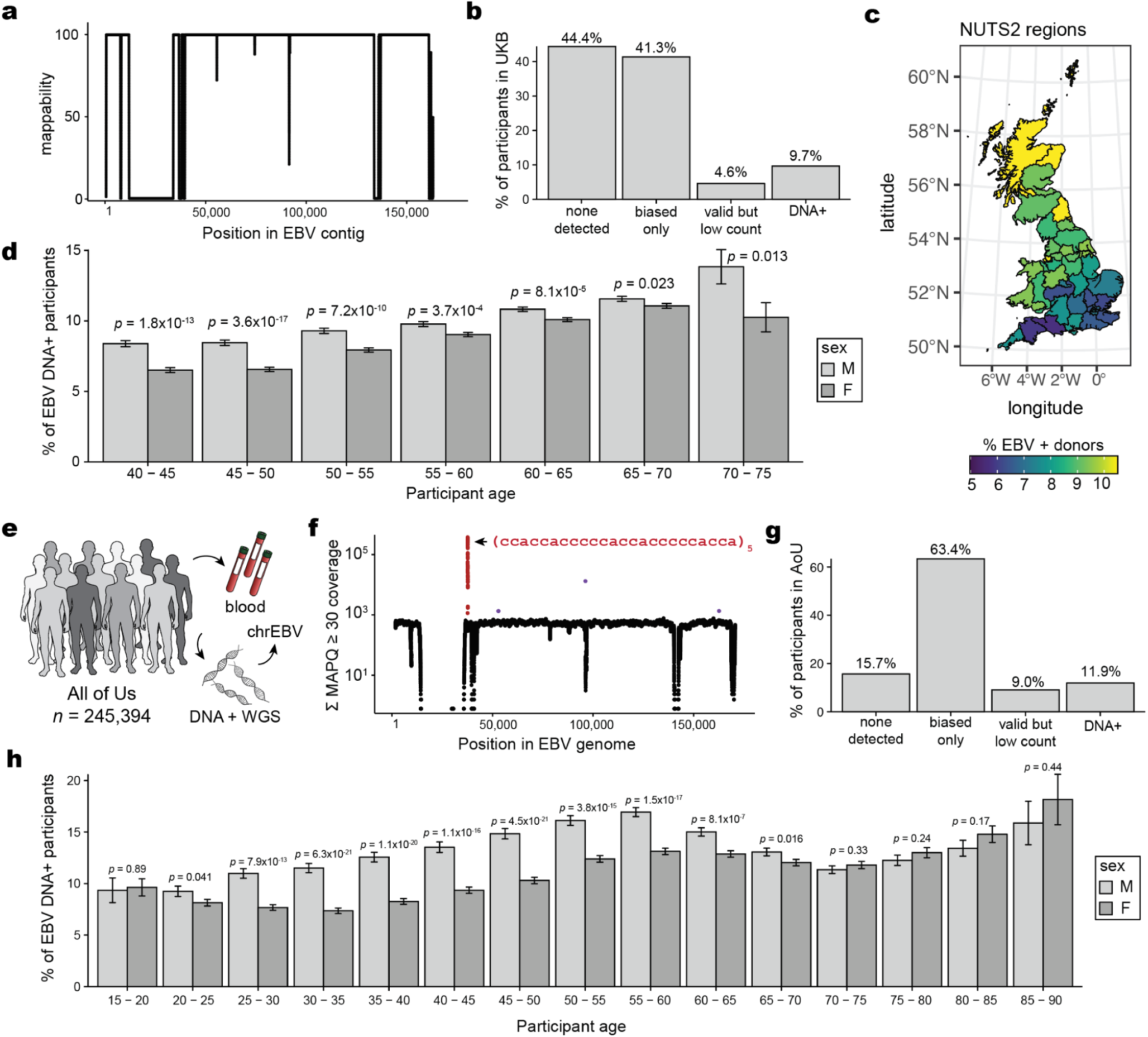
Supporting analyses of EBV DNA detected from WGS data. **(a)** Mappability of the EBV contig in the hg38 reference. **(b)** Partition of UKB participants by EBV DNA detection after accounting for biased regions. “Biased only” refers to participants with reads mapping to only the two repetitive regions indicated in Fig. 1b. “Valid and low count” refers to participants with EBV DNA detected after masking the two biased regions. “DNA+” refers to participants who pass the threshold of 1.2 EBV copies per 10^4^ human cells. **(c)** Geographical distribution of participant birth location colored by percent EBV DNA+, split by UK NUTS2 annotations. **(d)** Percent EBV DNA+ resolved by sex and age in UKB. Statistical test: two-sided proportion test comparing sex in associated age bin. Error bars: standard error of the mean. **(e)** Schematic of AoU chrEBV extraction from blood-based WGS. **(f)** Sum of per-base read coverage of map quality (MAPQ) score ≥30. **(g)** Same as b but for AoU. **(h)** Percent EBV DNA+ resolved by sex and age in AoU. Statistical test: two-sided proportion test. Error bars: standard error of the mean.

**Extended Data Figure 2.**
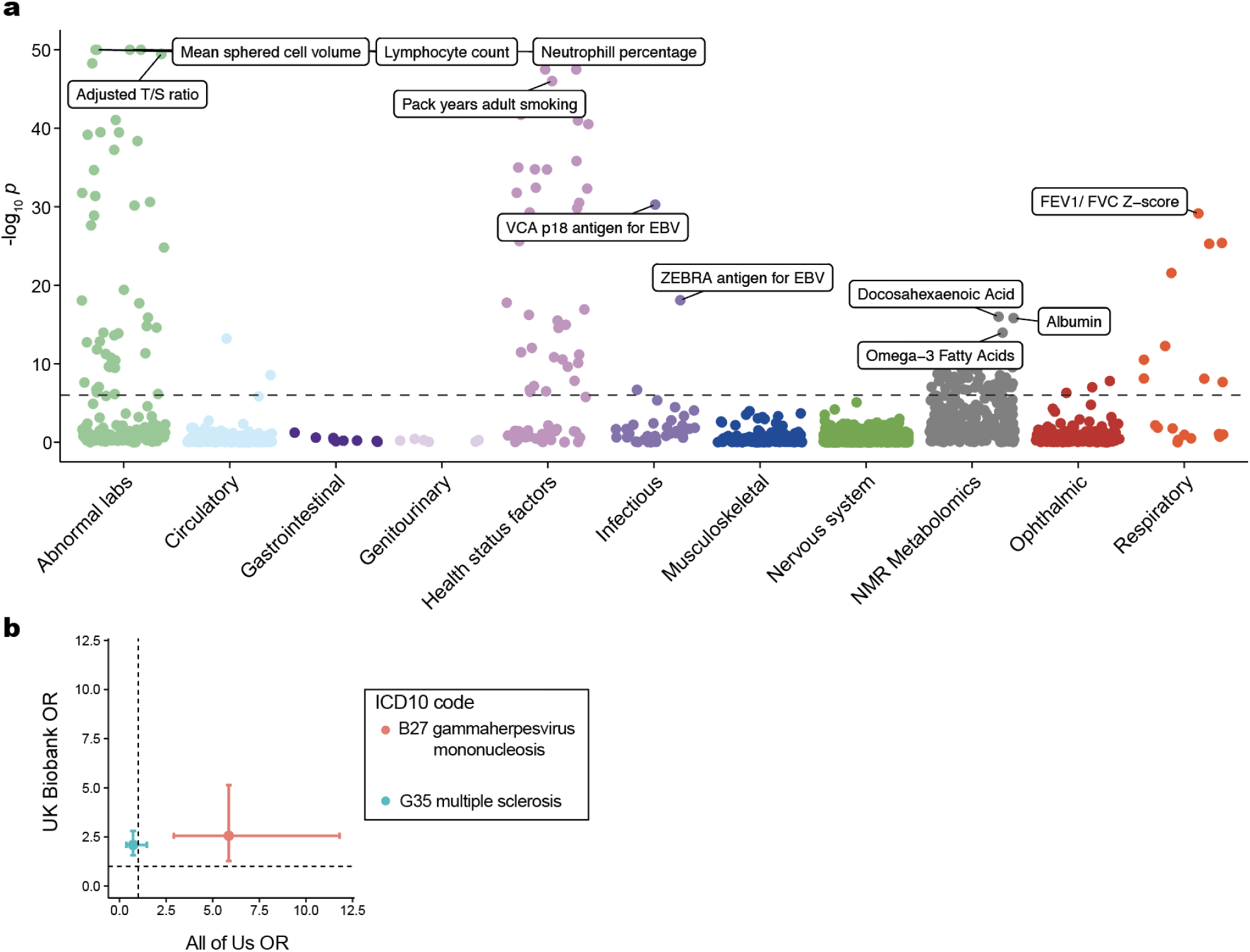
**Summary of phenome-wide associations for selected traits**. **(a)** Summary of associations between EBV DNAemia and quantitative traits in UKB. The dashed line represents the genome-wide significant *P* value threshold (3.3 × 10^-6^). The y-axis is capped at-log_10_(*P*) = 50; all associations are plotted (*n* = 1,931), and selected traits are highlighted based on biological interest. **(b)** Focused association summary for two ICD10 codes. Error bars represent the 95% confidence interval of the OR estimate from either cohort. Dotted lines at OR = 1 represent null associations.

**Extended Data Figure 3.**
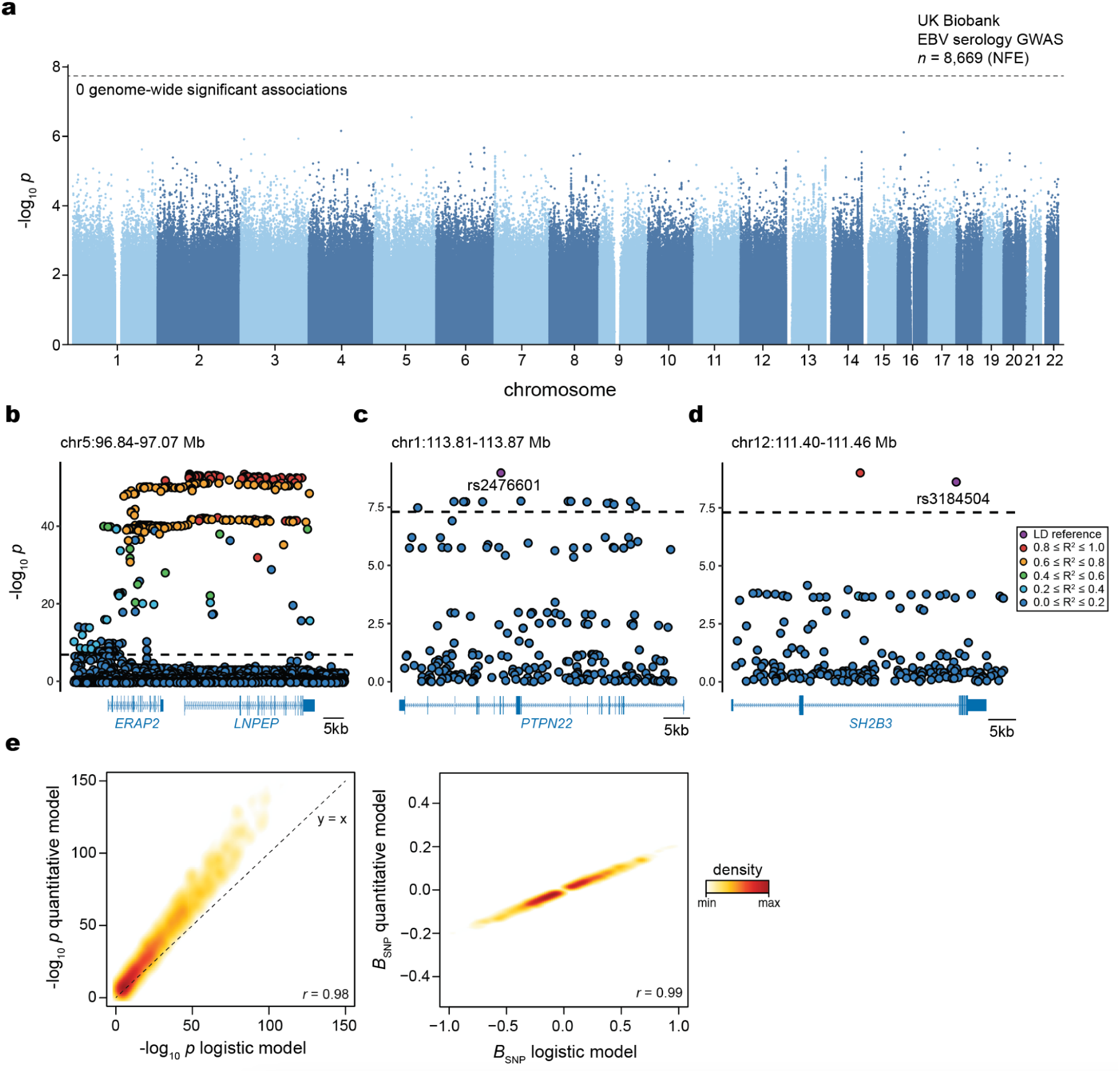
Supporting analyses for genetic association studies. **(a)** Manhattan plot of binarized EBV serostatus among 8,669 individuals of non-Finnish European (NFE) ancestry in UKB. **(b)** Zoom plot of the *ERAP2-LNPEP* locus in the EBV DNAemia GWAS with the UKB NFE cohort. **(c)** Zoom plot of the *PTPN22* locus in the EBV DNAemia GWAS with the UKB NFE cohort, highlighting the rs2476601 variant. **(d)** Zoom plot of the *SH2B3* locus in the EBV DNAemia GWAS with the UKB NFE cohort, highlighting the rs3184504 variant. **(e)** Comparison of EBV GWAS analyses as a quantitative trait (y-axis) compared to a logistic regression binarization (x-axis) for all variants *P* < 1× 10^-6^ in the EBV DNAemia GWAS with the AoU EUR cohort.-log_10_ p-values (left) and effect sizes (right) are shown at the variant level.

**Extended Data Figure 4.**
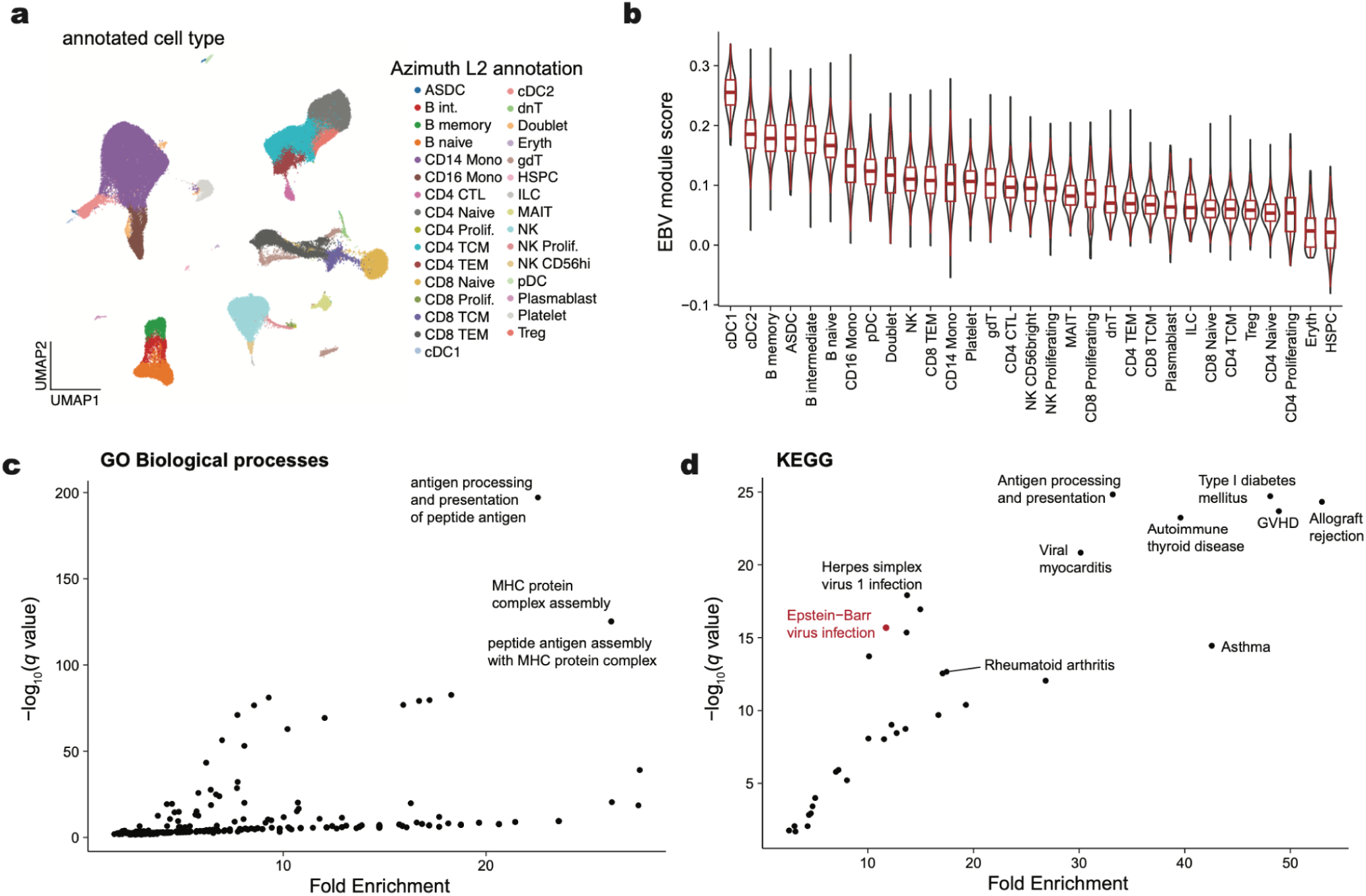
Supporting analyses of EBV DNAemia-associated genes at single-cell and pathway resolution. **(a)** UMAP embedding of 211,000 peripheral blood mononuclear cells. The broad cell type annotation (Azimuth L2^73^) annotates refined cell types. **(b)** Summary of EBV signature stratified by Azimuth L2 cell type, sorted by median score. Boxplots: center line, median; box limits, first and third quartiles; whiskers, 1.5× interquartile range. **(c)** Summary of GO Biological Processes and **(d)** KEGG gene set analyses. Top pathways based on *q-*value and fold enrichment are annotated.

**Extended Data Figure 5.**
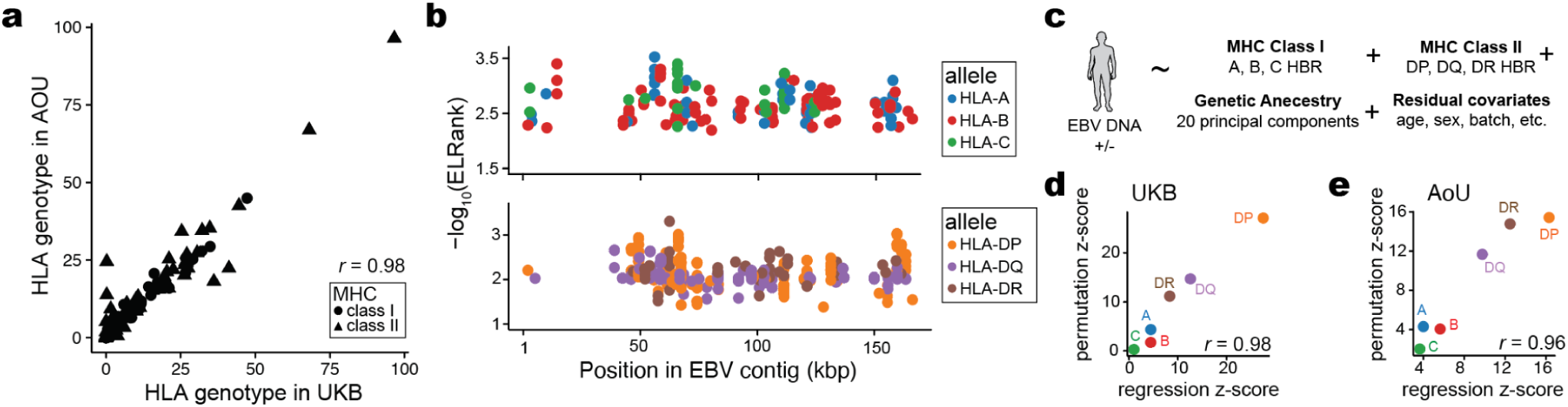
Supporting analyses of HLA-specific and antigen-binding associations. **(a)** Summary of four-digit HLA genotype frequencies across both biobanks. The Pearson correlation of the allele frequency is noted. **(b)** Annotation of the strongest predicted peptide among variable HLA alleles for class I (top) and class II (bottom). **(c)** Schematic of logistic regression analyses. **(d)** Correspondence between z-scores per allele for regression and permutation statistical models for the UKB cohort. The Pearson correlation between z-scores of the alleles is noted. **(e)** Same as in (d) but for the All of Us cohort.

**Extended Data Figure 6.**
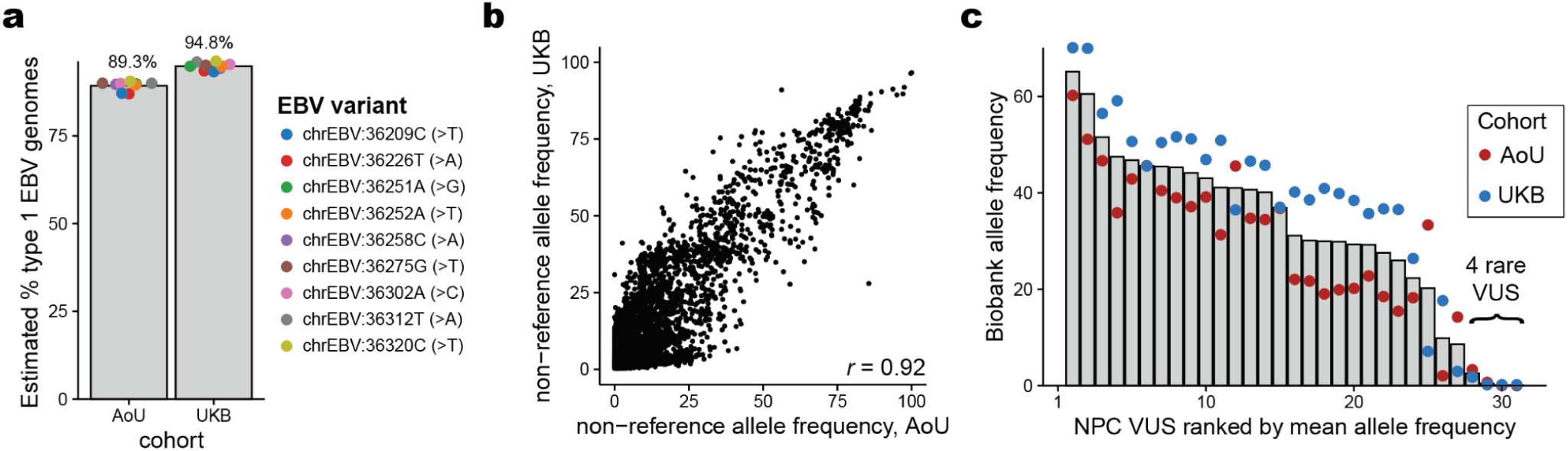
Analyses of genetic variation in the EBV genome. **(a)** Summary of 9 selected variants that discriminate between type 1 and type 2 EBV strains. The observed allele frequency for the reference contig (NC_007605; type I EBV) is plotted, and the corresponding type 2 allele is noted in parentheses. **(b)** The Pearson correlation of the two allele frequencies is noted. **(c)** Characterization of EBV variants of unknown significance (VUS) from cohorts of nasopharyngeal carcinoma (NPC) tumors^70^. All but four variants were detected at 10%+ in one or both cohorts.

